# Novel role of AGT gene in aplastic anaemia among paediatric patients based on gene expression profiling

**DOI:** 10.1101/2020.05.29.122861

**Authors:** Sarmistha Adhikari, Paramita Mandal

## Abstract

**Objectives:** Severe aplastic anemia is characterized by a hypocellular bone marrow and peripheral cytopenia. Mesenchymal stem cells (MSCs) play a crucial role in haematopoietic stem cells (HSCs) development and the microenvironment suitable for haematopoiesis. Investigation of the therapeutic targets by paediatric patient-specific gene expression analysis of the MSCs can be important for diagnosis.

**Methods:** The study was based on freely available miRNA and host gene expression in NCBI GEO dataset. Microarray based gene expression profiles (GSE33812) of MSCs for five paediatric aplastic anaemia patients and healthy controls were generated using Agilent-014850 platform and the data was downloaded from the database.

**Results:** MSCs gene expression profiling distinguished between healthy controls, children with aplastic anemia. Angioteninogen (AGT) gene involved in ERK1/ERK2 cascade, cyotokine secretion, metabolic processes was strongly down-regulated among all the patients with aplastic anemia. Emerging role of various transcription factors binding to this gene was identified as a new avenue of therapeutic application.

**Conclusions:** As a potential diagnostic tool, patient-specific gene expression profiling of MSCs made it possible to make the difficult diagnosis of most patients with aplastic anemia.

## INTRODUCTION

Aplastic anemia (AA) is a rare, immune-mediated hematopoietic disorder associated with significant morbidity and mortality [1]. AA can be diagnosed in patients presenting with pancytopenia and a hypocellular bone marrow. Typical symptoms include fatigue and easy bruising or bleeding; infections may be present, but generally there is no long-standing illness [2]. In patients with suspected AA, rapid and accurate diagnosis and concomitant supportive care are critical. Historically, immunosuppressive therapy (IST) and bone marrow transplantation (BMT) in eligible patients have been the mainstay of AA treatment [1]. In pediatric patients, new transplant strategies and improvements in supportive care have led to greatly improved outcomes and increasing use of BMT in both upfront and refractory settings [1, 3].

The incidence of AA varies with geography and it was found to be higher in Asia and lower in Europe, North America and Brazil according to the International Agranulocytosis and Aplastic Anemia Study [IAAAS] [4-7]. It was also identified that the incidence of that disease was 2-to 3-fold higher in Asia than in the West [8]. The great variation of the incidence of the disease is due to differential environmental exposure such as use of certain drugs and chemicals or by infectious agents such as viruses and bacteria. Besides the environmental agents the genetic background of different ethnic population may confer the risk of that disease [9-11]. It is really complicated to characterize the paediatric patients with aplastic anaemia than an adult because numerous inherited bone marrow failures can also present with aplastic anaemia without any obvious somatic features. Therfore, a precise diagnostic technique is essential for the children for the application of therapeutics [12]. Different demographic factors were already reported to be associated with aplastic anaemia among pediatric individuals and disease severity [13]. As the children are more sensitive to newer therapeutic agents in respect to their tolerability and suitability in contrast with chemotherapy or stem cell transplant, it’s necessary to establish and then incorporate into optimal treatment strategies.

Transcriptome analysis can clearly differentiate healthy controls from samples of AA patients. A study on transcriptome analysis among paediatric aplastic anaemia patients identified differentially expressed genes are involved in cell metabolism and cell communication or adhesion [14]. Fischer et al., 2012 able to identify that the transcription of major integrins was dramatically down-regulated in the few remaining CD34^+^ bone marrow cells from children with severe aplastic anaemia [15]. MSCs usually resides within the stroma and they are derived from bone marrow. They play significant role in hematopoiesis and immunomodulation. Various studies have reported the differential gene expression in MSCs among patients with aplastic anaemia compared to healthy controls. A study by Li et al., 2012 identified over 300 differentially expressed genes among aplastic anaemia compared with healthy controls [16]. These differentially expressed gene are involved in apoptosis, adipogenesis, and the immune response. Another study also reported increased MSC apoptosis in AA patients [17]. Consequent studies also revealed that MSCs among AA patients have lower proliferation potential [18, 19].

Various studies have already reported gene expression profiling of bone marrow MSCs from aplastic anaemia patients and they identified several genes those are involved in various biological processes such as cell cycle, cell division, proliferation, chemotaxis, adipogenesis-cytokine signalling and haematopoietic cell lineage differentiation which suggests that impaired cellular function is a hallmark of this disease [20-22]. It is well documented in several studies that transcription factor deregulation were also involved in aplastic anaemia. It was reported that GATA2 transcription factor downregulation in bone marrow MSCs can accelerate adipocyte differentiation which is one of the chief characteristics of aplastic anaemia [23].

Therefore, we undertook the current study to determine the deregulated genes associated with pediatric patients with aplastic anaemia and their transcription factor binding motifs which can regulate this disease progression.

## Materials and methods

### Source data

The expression analysis of the microarray datasets from the Gene Expression Omnibus [GEO] database of the National Center for Biotechnology Information [NCBI] of the U.S. National Library of Medicine was used for the current study. There was single gene expression profiling dataset for aplastic anemia patients. This dataset included 5 severe pediatric aplastic anemia patients.

### Defination of clinical dataset

For gene expression profiling, we have used GEO dataset with accession id GSE33812 [24] which included bone marrow mesenchymal stem cells of 5 pediatric aplastic anemia patients and also healthy donors. Agilent-014850 Whole Human Genome Microarray 4×44K G4112F GeneChips were used for gene expression profiling.

### Microarray based gene expression analysis

Gene expression profiles for all the categories of aplastic anaemia patients and healthy controls were generated using Agilent-014850 Whole Human Genome Microarray 4×44K G4112F probes. The probe quality of the array was assessed before and after normalization and the background correction was done using bioconductor based limma package. To improve data quality, a filtering of the probes was applied. The probes containing repetitive sequences, binding to multiple sites of human transcriptome, were removed for further analysis. Downstream analysis was done to identify the differentially expressed genes, based on t-test among both the categories of pediatric aplastic anemia patients compared to healthy controls. The p-values were determined and multiple testing corrections [Benjamini Hochberg method] done to remove the false discovery rate. The differentially expressed genes were selected on the basis of p□value<□0.05 and fold change of gene expression, compared to controls [for up-regulation, fold change ≥2 and for down-regulation, fold change ≤−2].

### Identification of biological processes of significantly enrichsed genes

To get a global view of relevant biological pathways associated with disease progression, we used GeneCodis 3 [25, 26, 27] and we considered significant p value (<0.05) for identification of relevant biological processes.

### Cellular localization of deregulated genes

To know about the cellular localization of the deregulated genes, we used GeneCodis 3 and we considered significant p value (<0.05) for identification of relevant motifs for transcription factors.

### Identification of transcription factors binding motif of significantly enriched genes

To get a global view of transcription factor binding sites of the deregulated genes associated with disease progression, we used GeneCodis 3 and we considered significant p value (<0.05) for identification of relevant motifs for transcription factors.

## Results

### Differentially expressed genes in microarray based gene expression profiling

To identify the differentially expressed genes in aplastic anaemia patients we compared the gene expression profiling of pediatric aplastic anaemia patients compared to healthy controls. The gene were selected as differentially expressed on the basis of p value (<0.05) and log fold change (>2 or <2). Microarray based global gene expression revealed that there were 24 genes that are differentially expressed in case of patients samples rather than control samples. These genes are down-regulated in aplastic anaemia samples compared to healthy donors. The expression pattern of these gene are depicted in **Table 1**. Among the entire significantly altered gene, angiotensinogen (AGT) was consistently downregulated among all the samples.

**Table 1.**
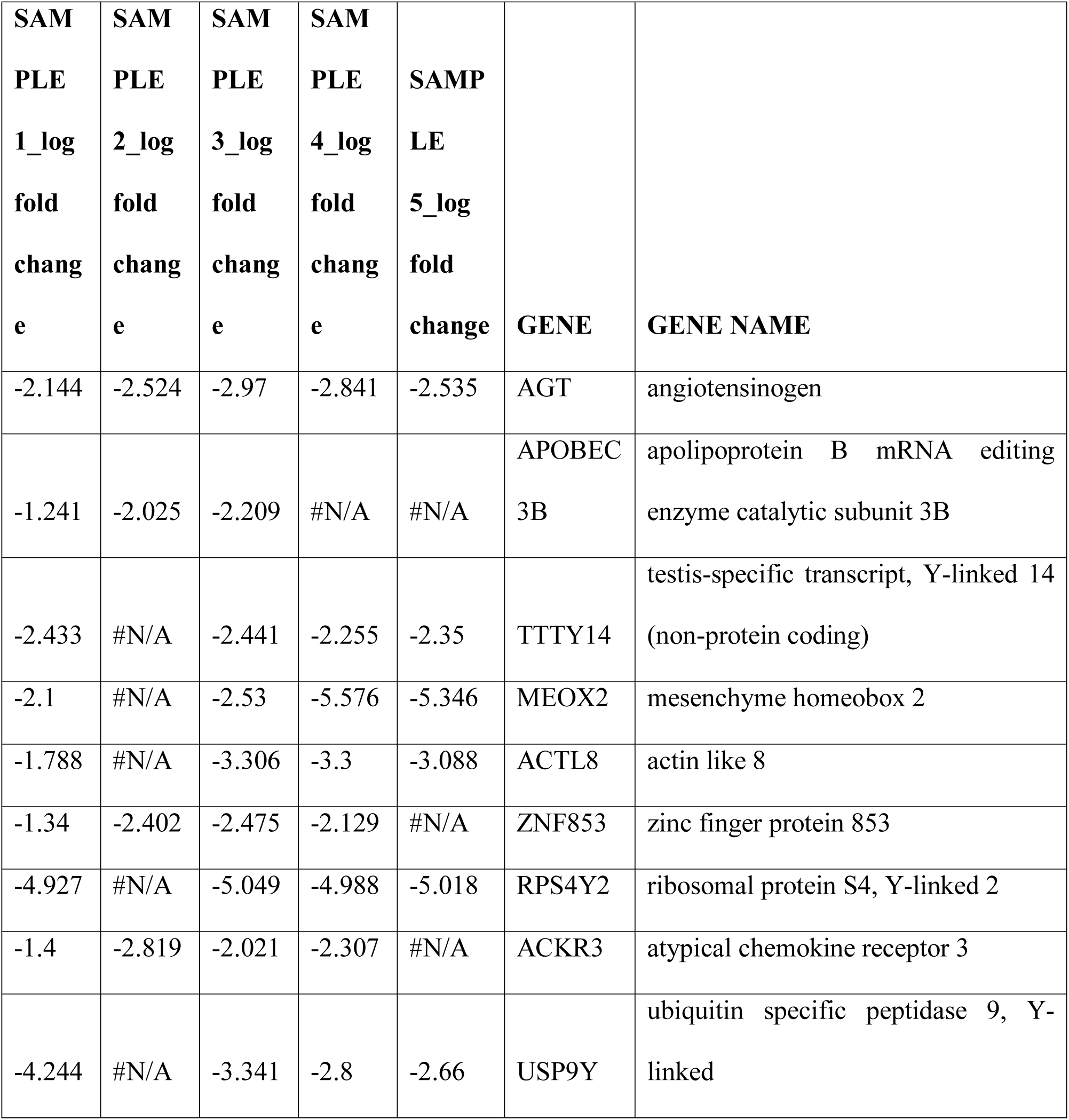
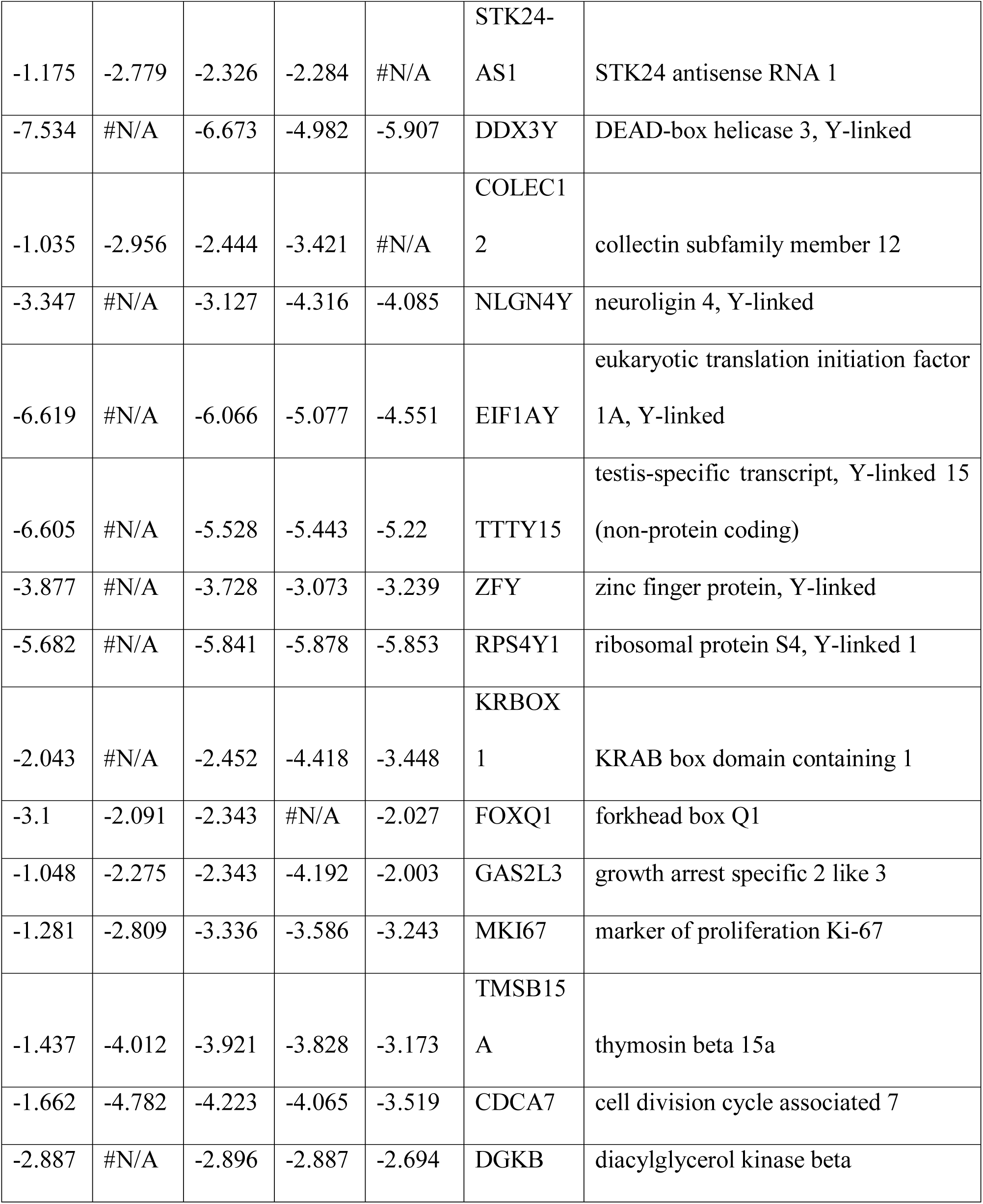
Significantly downregulated genes among paediatric patients with aplastic anaemia.

### Biological processes of differentially expressed genes

GeneCodis3 analysis of the differentially downregulated genes revealed that various cell signalling pathway, cell proliferation, metabolic pathways, translation, inflammatory and cytokine secretary pathways are significantly altered pathway among paediatric aplastic anaemia cases compared to healthy controls. The significantly altered downregulated genes in those pathways are AGT, FOXQ1, RPS4Y1, RPS4Y2, DGKB and CDCA7. The details of biological processes with the list of genes are depicted in **Table 2**.

**Table 2.** Significantly enriched biological processes among paediatric patients with aplastic anaemia.

| **Biological processes** | **p value** | **Genes** |
| --- | --- | --- |
| regulation of cell proliferation | 0.015<br>8403 | CDCA7,A<br>GT |
| cytokine secretion | 0.015<br>8892 | AGT |
| positive regulation of activation of JAK2 kinase activity | 0.016<br>9114 | AGT |
| G-protein signaling, coupled to cGMP nucleotide second messenger | 0.017<br>1986 | AGT |
| positive regulation of protein tyrosine kinase activity | 0.020<br>8347 | AGT |
| positive regulation of cellular protein metabolic process | 0.020<br>8347 | AGT |
| positive regulation of cytokine production | 0.023<br>0759 | AGT |
| translation | 0.023<br>7124 | RPS4Y2,<br>RPS4Y1 |
| activation of NF-kappaB-inducing kinase activity | 0.024 | AGT |
|  | 7379 |  |
| ERK1 and ERK2 cascade | 0.026<br>2278 | AGT |
| DNA fragmentation involved in apoptotic nuclear change | 0.030<br>4993 | FOXQ1 |
| activation of phospholipase C activity by G-protein coupled receptor<br>protein signaling pathway coupled to IP3 second messenger | 0.035<br>442 | AGT |
| phosphorylation | 0.039<br>9754 | DGKB |
| activation of protein kinase C activity by G-protein coupled receptor<br>protein signaling pathway | 0.039<br>9792 | DGKB |
| positive regulation of phosphatidylinositol 3-kinase cascade | 0.039<br>9821 | AGT |
| positive regulation of inflammatory response | 0.039<br>9821 | AGT |

### Identification of localization of differentially expressed gene

GeneCodis3 analysis of the differentially downregulated genes revealed the cellular localization of the significantly downregulated genes. Intracellular localization of RPS4Y2, MKI67, RPS4Y1 genes were rcorded. Moreover, cytoplasmic expression of DGKB, MKI67 and ribosomal localization of RPS4Y1 and RPS4Y2 are recorded. The details of cellular localization of those genes are depicted in **Table 3**.

**Table 3.**
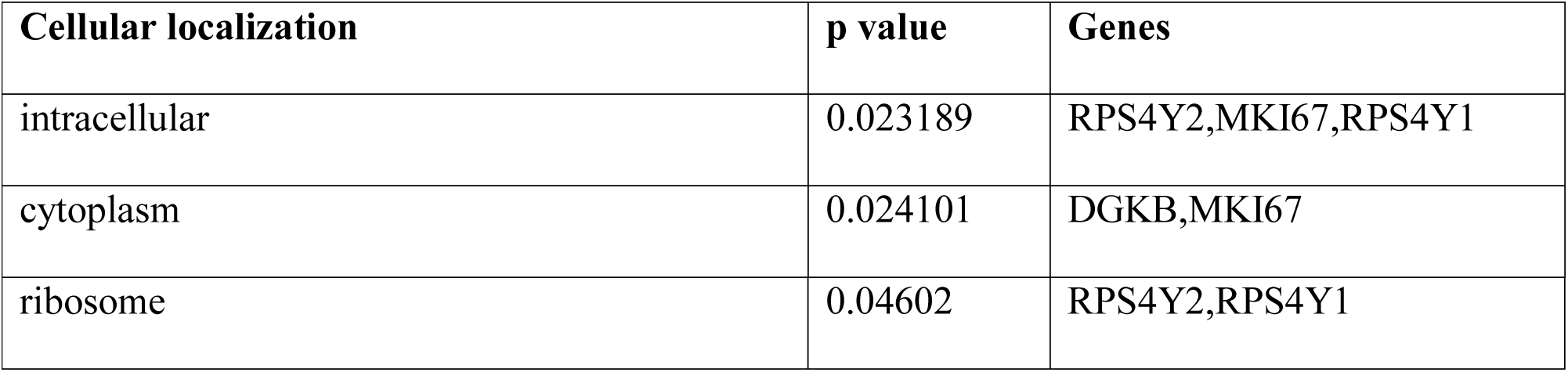
Cellular localization of significantly enriched genes among paediatric patients with aplastic anaemia.

### Determination of transcription factor binding sites of the differentially expressed genes

GeneCodis3 analysis of the differentially downregulated genes revealed some transcription factor binding site motifs of the significantly altered genes. This includes forkhead associated transcription factors, diacylglycerol kinase binding factors and ribosomal proteins. The details of transcription factor binding sites are depicted in **Table 4**.

**Table 4.** Transcription factor binding sites of significantly enriched genes among paediatric patients with aplastic anaemia.

| Transcription factor binding site | p value | Genes |
| --- | --- | --- |
| Forkhead-associated (FHA) domain | 0.02199 | MKI67 |
| Transcription factor, fork head | 0.0277 | FOXQ1 |
| Serpin domain | 0.019697 | AGT |
| Diacylglycerol kinase, accessory domain | 0.00525 | DGKB |
| Diacylglycerol kinase, catalytic domain | 0.007575 | DGKB |
| RNA-binding S4 | 3.24E-06 | RPS4Y2,RPS4Y1 |
| Ribosomal protein S4e | 1.95E-06 | RPS4Y2,RPS4Y1 |
| Angiotensinogen | 0.000585 | AGT |

## Discussion

To know about the biology of aplastic anaemia, we should know about the biology of mesenchymal stem cells which enable us to know about the bone marrow microenvironment. This enable us to manipulate human cells and we will be moving into an exciting phase of personalized herapy for bone marrow failure. Till date there was no such microarray based gene expression profiling of mesenchymal stem cells among paediatric aplastic anaemia patients. The microarray based gene expression profiling of mesenchymal stem cells of aplastic anaemia revealed 24 differentially downregulated genes among those patients compared to healthy donors.

Our study revealed that AGT gene was significantly downregulated among all the patients with aplastic anaemia. Angiotensinogen (AGT) is the precursor of the vaso-active peptide, angiotensin II. AGT gene moleculer variant was identified as a significant risk factor and hereditary marker of hypertension [28] which is also associated with cytokine secretion may be related to aplastic anaemia.

In our analysis we have found that ERK1 and ERK2 cascade, PI3K, cell proliferation, cytokine secretion are significantly downregulated among children with aplastic anaemia. It was already reported that ERK activity can be regulated in response to of hematopoietic cytokines and growth factors and it play critical roles in hematopoiesis [29] which justifies our finding. Mesenchymal stem cells play a vital role in haematopoeitic stem cell proliferation; they also modulate immune responses and maintain an environment supportive of hematopoiesis [30]. It is also well established that cytokine mediate hematopoietic stem cell development[31] and thus downregulation could be relevant for aplastic anaemia. Thus downregulation of cell proliferation could be relevant for aplastic anaemia.

Advanced knowledge on the transcription factors, in terms of structure, function (expression, degradation, interaction with co-factors and other proteins) and the dynamics of their mode of binding to DNA paved the way for new therapies targeted against transcription factors [32]. In this study we were able to identify various transcription factors (Forkhead-associated domain, fork head transcription factor, serpin domain, diacylglycerol kinase accessory domain, diacylglycerol kinase catalytic domain, RNA binding S4, ribosomal protein S4e, angiotensin) associated with altered gene expression which is really important for the therapeutic application.

Children with aplastic anaemia have promising therapeutic outcomes if treated adequately with antibiotics and referred early. Once the genetic nature is confirmed, novel drugs like small molecule inhibitors can be applied and offer hope for the future in the management of these children [33, 34]. Recent study from our lab showed the potential of miRNA as therapeutic target for aplastic anaemia [35]. Thus it may be a indication that this particular transcription factor or drug may be helpful to use as a therapeutic agent in upcoming days. So, if we alter the particular gene expression and use transcription factor and novel drug as a therapeutic tool in case of paediatric aplastic anemia patients, this could be helpful for those patients suffering from aplastic anemia.

**Figure 1:**
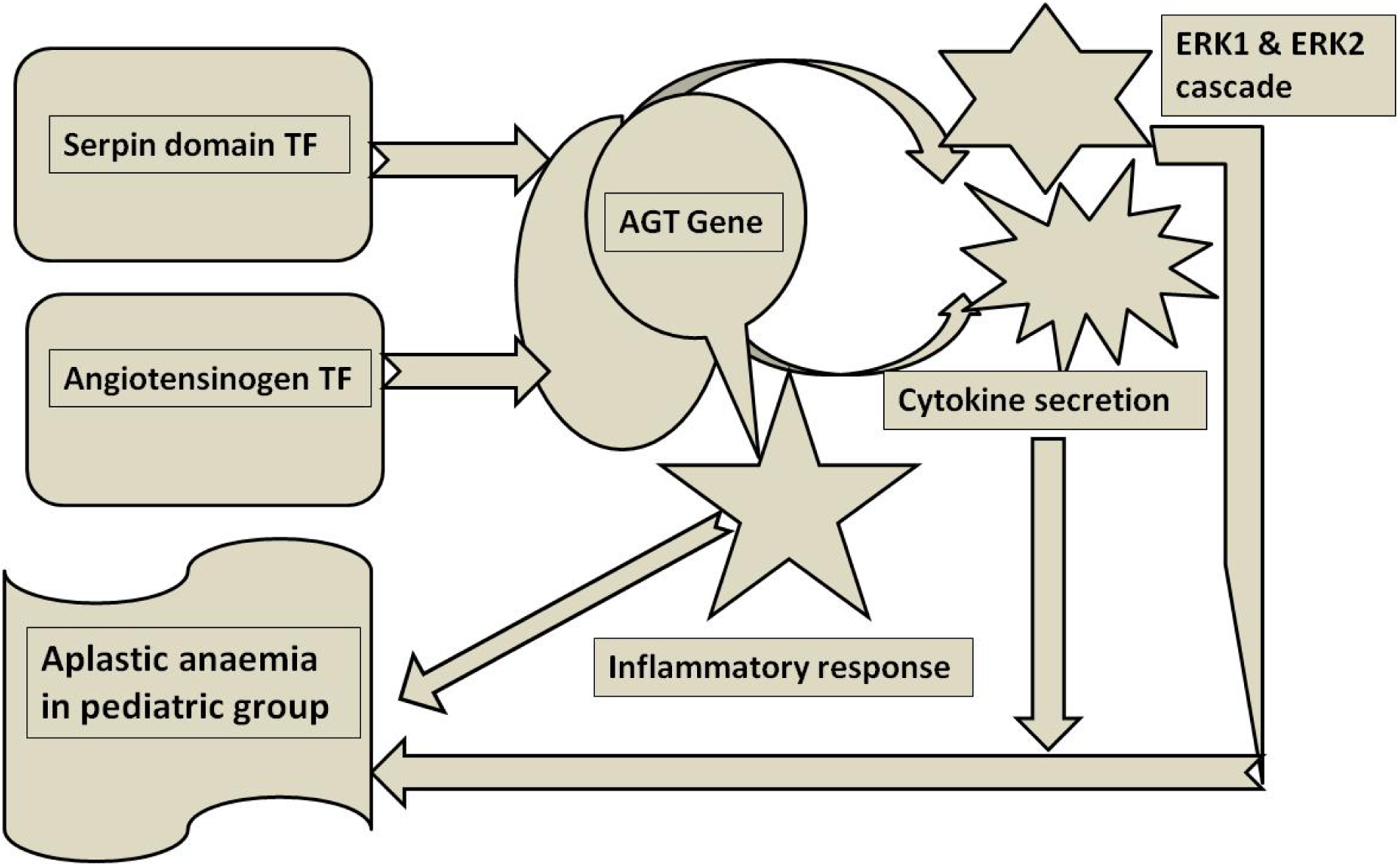
Schematic diagram of the role of AGT gene in aplastic anaemia among paediatric patients.

## Acknowledgement

We thank all the clinicians and researchers who performed the experiments and made accessible online datasets. We thank Chow K et al. of National Chung Hsing University, Taiwan for doing gene expression profiling of paediatric aplitic anaemia patients and healthy donors and made it accessible in GEO database. The financial support for the study was provided by Department of Biotechnology, Govt. of India [Grant id: BT/PR18640/BIC/101/924/2016 DATED 20.09.2017]. Last but not the least; we are thankful to the Department of Zoology, The University of Burdwan (DST-FIST and PURSE) for the infrastructural support.

SA and PM: Conceived the objectives and performed the analysis, SA and PM: wrote the paper.

## Ethical approval

This article does not contain any studies with human participants performed by any of the authors.

